# Galanin inhibits visceral afferent responses to noxious mechanical and inflammatory stimuli

**DOI:** 10.1101/746586

**Authors:** Toni S. Taylor, Parvesh Konda, Sarah S. John, David C. Bulmer, James R.F. Hockley, Ewan St. John Smith

**Affiliations:** Department of Pharmacology, University of Cambridge, Tennis Court Road, Cambridge CB1 2PD, UK

**Author notes:** Corresponding author: Ewan St. John Smith PhD, Department of Pharmacology, University of Cambridge, Tennis Court Road, Cambridge CB1 2PD, UK. GSK, GSK Medicines Research Centre, Gunnels Wood Road, Stevenage, Hertfordshire, SG1 2NY, UK.

**Keywords:** Colon, Visceral pain, Galanin, Inflammation, Mechanosensitivity, Hypersensitivity

## Abstract

Galanin is a neuropeptide expressed by sensory neurones innervating the gastrointestinal (GI) tract. Galanin displays inhibitory effects on vagal afferent signalling within the upper GI tract, and the goal of this study was to determine the actions of galanin on colonic spinal afferent function. Specifically, we sought to evaluate the effect of galanin on lumbar splanchnic nerve (LSN) mechanosensitivity to noxious distending pressures and the development of hypersensitivity in the presence of inflammatory stimuli and colitis. Using *ex vivo* electrophysiological recordings we show that galanin produces a dose-dependent inhibition of colonic LSN responses to mechanical stimuli and prevents the development of hypersensitivity to acutely administered inflammatory mediators. Using galanin receptor (GalR) agonists, we show that GalR1 activation, but not GalR2/3 activation, inhibits mechanosensitivity. The inhibitory effect of galanin on colonic afferent activity was not observed in tissue from mice with dextran sodium sulphate-induced colitis. We conclude that galanin has a marked inhibitory effect on colonic mechanosensitivity at noxious distending pressures and prevents the acute development of mechanical hypersensitivity to inflammatory mediator, an effect not seen in the inflamed colon. These actions highlight a potential role for galanin in the regulation of mechanical nociception from the bowel and the therapeutic potential of targeting galaninergic signalling to treat visceral hypersensitivity.

**Key point summary:**

- Galanin inhibits visceral afferent mechanosensitivity to noxious phasic distension of the colon via GalR1.
- Galanin attenuates afferent mechanical hypersensitivity induced by the application of inflammatory mediators.
- Inhibition of afferent mechanosensitivity by galanin is not observed in tissues isolated from mice undergoing DSS-induced colitis
- Galanin inhibits the transmission of noxious mechanical stimuli by colonic afferents and its sensitisation by inflammatory mediators highlighting an antinociceptive role for galanin in the colon.

## Introduction

Visceral hypersensitivity is commonly observed in patients with gastrointestinal (GI) diseases, such as irritable bowel syndrome (IBS) and inflammatory bowel disease (IBD), and contributes to the presence of pain in these conditions (Chang et al., 2000; Ritchie, 1973). In the periphery, distal terminals of sensory neurones transduce noxious stimuli to produce nociception and the sensation of pain (Grundy et al., 2019; Hockley et al., 2018; St John Smith, 2018). The persistent activation of these sensory nerve endings in response to disease driven changes in the bowel microenvironment (e.g. the production of inflammatory mediators such as prostaglandins), coupled with longer term changes in the plasticity of the sensory nerve endings and subsequent sensitisation of central pain pathways, contribute to the development of visceral hypersensitivity and represent potential targets for therapeutic intervention. The distal colon receives dual sensory innervation from afferent fibres running within the lumbar splanchnic nerve (LSN) and the pelvic nerve (PN) (Deiteren et al., 2015). Both nerves have been implicated in the processing of pain from the colorectum, with the PN being identified as the predominant pathway for the signalling of pain in response to rectal distension in humans, and the LSN being identified as the predominant pathway for the signalling of pain in responses to distension of the sigmoid and descending colon (Hughes et al., 2009; Brierley et al., 2004; Ray and Neill, 1947). Therefore, the modulation of LSN and PN signalling from respective colonic and rectal regions have important implications in the treatment of chronic visceral pain in GI diseases such as IBS and IBD (Brierley et al., 2004; Siri et al., 2019).

One method by which colorectal sensory nerves can be modulated is through local release of peptides, such as calcitonin gene-related peptide (CGRP), contained within the nerves themselves, or from non-neuronal cells within the GI tract (Lakhan and Kirchgessner, 2010). We have shown through single-cell RNA sequencing that the peptide galanin is highly expressed in colonic sensory neurones that also express Trpv1, i.e. putative nociceptors, suggesting a potential role in the regulation of colonic nociception (Hockley et al., 2019). Galanin acts upon 3 receptors, GalR1, GalR2 and GalR3 (Branchek et al., 2000, 1998), and single-cell RNA sequencing demonstrates that GalR1 is the most highly expressed in colonic sensory neurones and that its expression, along with that of GalR2, is largely restricted to Trpv1 positive thoracolumbar DRG arising from the LSN, thus suggesting a paracrine role for galanin in the regulation of sensory signalling from the bowel (Hockley et al., 2019). When activated by galanin, GalR1 predominantly signals through G_i_, suggesting that the effects of galanin on colonic afferent signalling will be inhibitory (Branchek et al., 2000). Consistent with this supposition, in the upper GI tract of mice and ferrets, galanin produces inhibition of vagal afferent signalling through the activation of GalR1 receptors. Galanin has also been shown to have a stimulatory effect on vagal afferents through the activation of its G_q_ coupled GalR2 receptor, with GalR3 receptors making no clear functional contribution despite its expression in the relevant ganglia (Page et al., 2007, 2005).

The role of galaninergic signalling in the colorectum is currently unclear and in this study, we set out to determine the effect of galanin upon LSN mechanosensitivity due to the high expression of both galanin and its receptors in the relevant thoracolumbar sensory ganglia. Moreover, because single-cell RNA-sequencing data demonstrate coexpression of GalR1 with receptors for certain inflammatory mediators (e.g. bradykinin and 5-hydroxytryptamine) and ion channels such as Trpv1 that are modulated by inflammatory mediators (Hockley et al., 2019), we wanted to determine if galanin could inhibit mechanical hypersensitivity induced by inflammatory mediators and extend this to an *in vivo* model of acute colitis.

We find that galanin is expressed by putative nociceptors originating from the LSN and that it inhibits LSN mechanosensitivity via GalR1. Inflammatory mediator induced mechanical hypersensitivity is also abolished by galanin, but that inhibition of mechanical hypersensitivity is lost in LSN obtained from mice with colitis.

## Materials and Methods

### Ethical Approval

C57BL/6J mice were used for all experiments. All protocols were performed in accordance with the UK Animals (Scientific Procedures) Act 1986 Amendment Regulations 2012 following ethical review by the University of Cambridge Animal Welfare and Ethical Review Body.

### Ex vivo mouse LSN preparation and recording

Adult (8-25 weeks) C57BL/6J male and female mice (Envigo) were humanely killed by cervical dislocation and exsanguination, and the distal colon (from the splenic flexure to rectum) with associated LSN removed. Colonic content was flushed with Krebs buffer and the colon tied to either end of a cannula, and perfused luminally (200 μL min^-1^), and serosally (7 mL min^-1^) with carboxygenated (95% O_2_, 5% CO_2_) Krebs buffer, in mM: 124 NaCl, 4.8 KCl, 1.3 NaH_2_PO_4_, 2.5 CaCl_2_, 1.2 MgSO_4_·7H_2_O, 11.1 glucose, and 25 NaHCO_3_; supplemented with indomethacin (3 μM, to suppress prostanoid synthesis), and nifedipine (10 μM) and atropine (10 μM) to block smooth muscle contraction as previously described (Hockley et al., 2014). The bath was maintained at 32-34°C. The inferior and superior mesenteric ganglia were identified at the point of the iliac bifurcation and suction electrode recordings of multiunit activity were made from neurovascular bundles isolated central to the inferior mesenteric ganglia. Signals were amplified at a gain of 5K, band pass filtered (100-1300 Hz), digitally filtered for 50 Hz noise (Humbug, Quest Scientific, Canada) and data acquired at 20 kHz (micro1401; Cambridge Electronic Design, UK). Ongoing nerve discharge was quantified from spikes passing a threshold level set at twice the background noise (typically ∼100 μV). All signals were displayed on a PC using Spike 2 software. The baseline pressure was set up at 2-3 mmHg and recordings were maintained for approximately 30 minutes before initiating the experimental protocols.

Mechanosensitivity was evaluated using ramp and phasic distension protocols. For the ramp distension protocol the luminal outflow cannula was blocked and the subsequent increase in pressure was observed until the desired maximum of 80 mmHg was reached (∼2.5 minutes), at which point the luminal outflow was re-opened. For the phasic distension protocol a rapid increase in the intraluminal pressure from 0-80 mmHg was achieved by switching the luminal outflow to a water column of sufficient height.

For drug treatments, two protocols were used. In one protocol three phasic distensions were performed to obtain a stable response to distension, and then galanin or vehicle (or the GalR1/2 agonists M617 and spexin respectively, Tocris) were serosally superfused (20 mL volume) between the third and fourth phasic distension. In the second protocol the effect of galanin (or vehicle) was examined on ramp and phasic distension (an initial ramp distension followed by three phasic distensions) in non-sensitised preparations (continual luminal perfusion with Krebs buffer) or sensitised preparations. Sensitisation was achieved by intraluminal perfusion of an inflammatory soup that has previously been shown to increase resting afferent activity (Su and Gebhart, 1998: 10 μM histamine, 10 μM prostaglandin E_2_, 10μM 5HT, 1 μM bradykinin, and 1 mM ATP) for 20 minutes prior to and during subsequent ramp and phasic distensions.

### Analysis of electrophysiological recordings

Phasic distension of the colon by 80 mmHg leads to an increase in LSN activity, which consists of an initial peak response that reduces in magnitude to a sustained increase in nerve discharge for the remainder of the distension period. Nerve discharge returning to baseline levels following cessation of the phasic distension. Peak changes and time profiles of LSN activity were determined by subtracting baseline firing (average over 60 seconds before distension) from increases in LSN activity following distension. During phasic distensions, peak firing was defined as the maximal firing rate observed during distensions, occurring within the first 15 seconds, and sustained firing frequency was defined as the activity seen during the subsequent 45 seconds of the phasic distension. Drug effect was determined by comparing an average of the first 3 distensions to the response of the third post-drug distension. By contrast, ramp distension leads to a slow and steady increase in activity until the end of the distension. Pressure profiles of LSN activity were determined by subtracting baseline firing from peak LSN activity measured every 5 mmHg.

### Retrograde labelling of colonic sensory neurones

Retrograde labelling of colonic sensory neurones was conducted as previously described (Hockley et al., 2019). In brief, C57BL/6J mice were anaesthetized with isoflurane (4% induction and 1-2% maintenance) then a midline laparotomy (∼1.5 cm incision) performed to reveal the distal colon. Five injections of 0.2 μL Fast Blue (2% in saline, Polysciences GmbH, Germany) were made into the wall of the distal colon using a glass needle at a rate of 0.4 μL min^-1^ using a microinfusion pump (Harvard Apparatus). After the abdominal cavity was flushed with saline to remove any excess Fast Blue dye, the muscle and skin layers were sutured and secured using 4-6 Michel clips. Postoperative care and analgesia (buprenorphine 0.05-0.1 mg kg^-1^) was provided and a glucose enriched soft diet provided, with regular checks of body weight. After a minimum of three days, the animals were killed using sodium pentobarbital (200 mg kg^-1^ i.p.) and transcardially perfused with phosphate buffered saline (PBS) followed by paraformaldehyde (4% in PBS; pH 7.4). Dorsal root ganglia (DRG; T13 – L1) were removed and further fixed in 4% paraformaldehyde for 30 minutes at 4 °C before cryoprotection in 30% sucrose overnight at 4 °C. The tissue was then embedded in Shandon M-1 Embedding Matrix (Thermo Fisher Scientific), snap frozen in liquid nitrogen, and stored at -80 °C until needed. Cryostat (Leica, CM3000; Nussloch) sections (12 μm) were collated across 10 slides (Superfrost Plus, Thermo Fisher Scientific) for each DRG.

### Immunohistochemistry

DRG sections were washed with PBS (twice for 2 minutes) and then blocked using antibody diluent (10% donkey serum, 5% bovine serum albumin and 0.2% Triton X-100 in 0.1 M PBS) for one hour. This was followed by overnight incubation at 4 °C with the appropriate primary antibodies (Table 1, we thank Prof. Theodorsson, Linköping University, for the kind gift of the anti-galanin antibody). The sections were then washed three times for five minutes with PBS and then incubated for 2 hours at room temperature with the appropriate fluorophore conjugated secondary antibodies (Table 2). No labelling was observed in control experiments where the primary antibody was excluded or in the presence of galanin as a blocking peptide for the anti-galanin antibody.

**Table 1.** List of primary antibodies

| Primary Antibody | Conc. | Company | Catalogue # | RRID | Reference |
| --- | --- | --- | --- | --- | --- |
| Rabbit anti-galanin | 1:1000 | Theodorsson Lab | Kind gift | - | Theodorsson et al., 2002 |
| Guinea pig anti-TRPV1 | 1:1000 | Alomone Labs | AGP-118 | AB_2721813 | Sanz-Salvador et al., 2011 |
| Goat anti-Gfrα3 | 1:300 | R&D Systems | AF2645 | AB_2110295 | Albers et al., 2014 |
| Goat anti-CGRP | 1:500 | Abcam | AB36001 | AB_725807 | Bravo-Herandez et al., 2016 |

**Table 2.** List of secondary antibodies

| Secondary Antibody | Conc. | Company | Catalogue # | RRID |
| --- | --- | --- | --- | --- |
| donkey anti-rabbit IgG-Alexafluor-488 | 1:1000 | Invitrogen | A-21206 | AB_2535792 |
| donkey anti-rabbit IgG-Alexafluor-568 | 1:1000 | Invitrogen | A10042 | AB_2534017 |
| donkey anti-goat IgG-Alexafluor-568 | 1:1000 | Invitrogen | A-11057 | AB_2534104 |
| donkey anti-guinea pig IgG-Alexafluor-488 | 1:1000 | Jackson Immuno Research | 706-165-148 | AB_2340460 |

### Imaging and quantification

Sections were imaged using an Olympus microscope (BX51) with Qicam camera and the relative intensities of DRG neurones after immunostaining were measured (ImageJ 1.51n analysis software, NIH, USA). The mean background intensity was subtracted to control for variability in illumination between images. Percentages of relative intensities were determined by comparison with least intensely (0%) and the most intensely (100%) labelled cells for each section. Relative intensities are calculated by subtracting the relative intensity of the darkest neuronal profile (a) from the relative intensity of the cell of interest (b) and comparing this to the relative intensity of the brightest neuronal profile (c) with the relative intensity of the darkest neuronal profile subtracted: relative cell intensity = (b – a)/(c – a) (Fang et al., 2002); cells with intensity values greater than the mean intensity of the darkest neuronal profiles from all the sections plus five times its standard deviation (SD) were considered positively labelled.

### Dextran sulfate sodium (DSS) model of induced colitis

C57BL/6J mice of either sex (8-12 weeks old) were weighed two days prior to the procedure; weight and stool content/consistency were then monitored daily throughout the treatment. 3% DSS (40,000 MW, Alfa Aesar) supplemented drinking water was administered with control mice receiving the same drinking water without DSS. The mice received DSS treated water for 5 days after which it was replaced with normal drinking water for a further 2 days. The experimenter was blinded, solutions labelled A and B, being unblinded after results were fully analysed. Oral administration of DSS leads to weight loss, diarrhoea and blood in the stool, which was scored to produce a disease activity index (DAI) (Table 3) (Manicassamy and Manoharan, 2014). On day 7, mice were killed, and relevant tissue samples and measurements were obtained. An approximately 1 cm section of colon was removed about 4 cm from the anus and fixed in 4% paraformaldehyde for 4 hrs and cryoprotected in 30% sucrose overnight. The tissue was then embedded in O.C.T mounting media (VWR Q-path Chemicals), snap frozen in liquid nitrogen, and stored at -80 °C until needed.

**Table 3.** Disease activity index (DAI)

| Score | Weight Loss | Stool Consistency | Blood in Stool |
| --- | --- | --- | --- |
| 0 | None | Normal | Normal |
| 1 | 1 – 5 % |  |  |
| 2 | 5 – 10 % | Very Soft | Slight bleeding |
| 3 | 10 – 15 % |  |  |
| 4 | 15 % | Watery diarrhoea | Gross bleeding |

### Histology: H&E with alcian blue staining

Cryostat sections of colon (20 μm) were collected and stored at -20 °C until needed. Slides were washed in tap water for 2 minutes before staining for 5 minutes with haematoxylin (1:2 dilution with tap water; Sigma), washing in tap water for 3 minutes followed by 0.3% HCl in ethanol for 30 seconds and then immediately washing in tap water for a further 2 minutes, before incubating in tap water until the tissue developed a deep blue appearance. Slides were then stained with alcian blue (1% W/V in 3% acetic acid; Polysciences Inc) for 10 minutes and washed in tap water for 2 minutes before being immersed in 100% ethanol for 30 seconds. Slides were then stained with eosin (Acros Organics) for 90 seconds, washed in tap water for 1 minute and then dehydrated in 100% ethanol for 30 seconds followed by 70% ethanol for 30 seconds. Slides were then cleared using Histoclear (National Diagnostics) for 30 seconds before mounting coverslips with Mowiol mounting media. Imaging was carried out using a NanoZoomer S60 Digital slide scanner. A histopathological score was made based on presence of inflammatory cells, crypt damage and ulceration, visualised using H&E staining and alcian blue to stain the goblet cells in the crypts

### Biotinylated hyaluronan binding protein (HABP) staining

20 μm colon sections were cut using a cryostat and mounted on glass slides, which were washed twice with PBS-Tween before being blocked with an antibody diluent solution (0.2% Triton X-100 and 5% bovine serum albumin in PBS) for one hour at room temperature. Slides were incubated in biotinylated hyaluronan binding protein (Amsbio, 1:200) at 4 °C overnight and then washed three times in PBS-Tween and incubated for two hours at room temperature with Alexafluor 488 conjugated streptavidin (Invitrogen, 1:1000). Slides are then washed three more times with PBS-Tween before being mounted and imaged using an Olympus BX51 microscope and QImaging camera.

### Statistics

All statistical analyses were performed using GraphPad Prism 6. IC_50_ values were derived by a sigmoidal dose-response (variable slope) curve using GraphPad Prism software. For ramp distensions, a repeated measures two-way ANOVA with Sidak’s post-hoc test was used. Both basal firing and phasic distension data were analysed using a repeated measures one-way ANOVA with Tukey’s post-hoc test. For the validation of the DSS model, DSS data were compared to control data using Student’s t-test. Statistical significance was set at P < 0.05. Data are presented as mean ± SD, N = number of animals, and n = number of cells.

## Results

### Galanin inhibits mechanically evoked LSN firing

Single-cell RNA-sequencing of colonic sensory neurones suggests that galanin is expressed in putative nociceptors (Hockley et al., 2019) and we confirmed this through immunohistochemistry on TL DRG isolated from mice in which the distal colon had been injected with the retrograde tracer fast blue. Galanin labelled 22.7 ± 8.3% of colonic sensory neurones (N = 3, n =466) and 4.5 ± 2.5% of all TL sensory neurones (N = 3, n = 1516). When examining coexpression with Gfrα3, the expression of which correlates with Trpv1, a marker for high-threshold stretch sensitive colorectal afferents (Malin et al., 2009), we observed marked co-localisation between Gfrα3 and galanin in colonic sensory neurones: Gfrα3 staining was present in 90.6 ± 1.2 % of galanin positive TL colonic sensory neurones (N = 3, n = 71, Fig. 1A, left panel) compared with overall Gfrα3 expression of 43.3 ± 7.5% in TL colonic sensory neurones (N = 3, n = 407). Similarly, the nociceptive marker Trpv1 was also highly enriched in galanin positive TL colonic sensory neurones (97 ± 2.5 % N = 3, n = 48, Fig. 1A, middle panel) compared with overall expression of 48.7 ± 3.3% (N = 3, n = 380) of TL colonic sensory neurones. Lastly, galanin was highly coexpressed with the peptidergic neuronal marker CGRP in colonic sensory neurones: 86.5 ± 6.7 % of galanin positive TL colonic sensory neurones were also CGRP positive (N = 3, n = 47, Fig. 1A, right panel). These results align with published RNA-sequencing data (Hockley et al., 2019) and confirm that galanin is highly expressed in nociceptive colonic sensory neurones.

**Figure 1.**
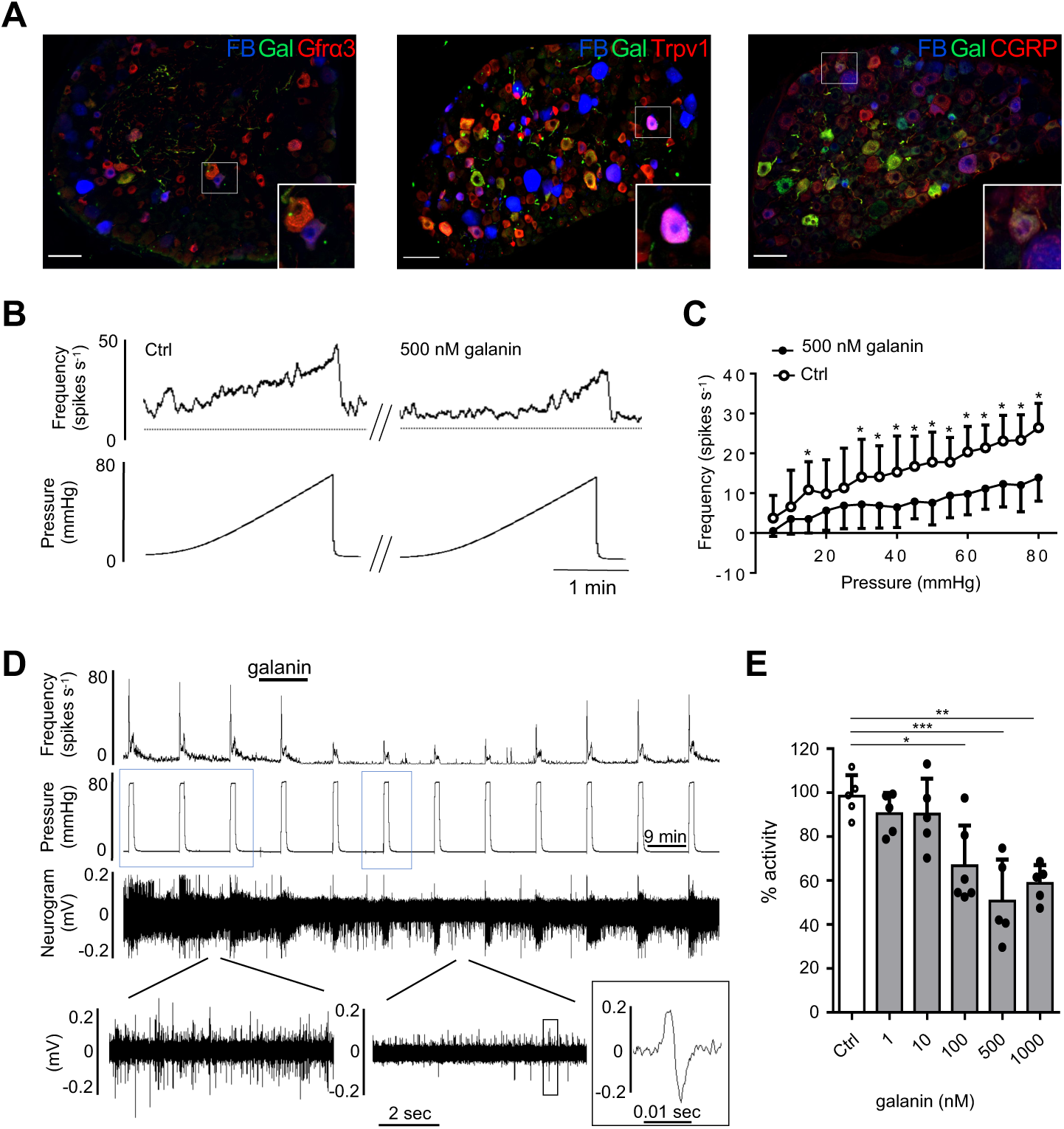
Galanin is expressed in colonic sensory neurones and inhibits LSN mechanosensitivity. (A) Galanin is expressed in TL colonic sensory afferents labelled with fast blue (FB) and is co- expressed with the nociceptive markers Gfrɑ3 (left) and Trpv1 (middle), and the peptidergic marker CGRP (right); scale bar 50 µm. (B) Example raw trace of ramp distension and response showing inhibition of activity by galanin. (C) Galanin inhibits the LSN response to ramp distension over a wide range of pressures (two-way repeated-measures ANOVA with Sidak’s post-hoc test, N = 6). (D) Example raw trace and frequency histogram of the LSN response to repeated phasic intraluminal distension (0 to 80 mmHg), showing reversible inhibition by galanin. Below, expanded sections showing basal activity before (left) and after (right) galanin application and a single action potential (black box). Blue boxes indicating distensions used for analysis. (E) Galanin dose dependently inhibited the peak response to phasic distension (one-way ANOVA with Tukey’s test, N = 6 for 100 nM and N = 5 for all other groups). Firing frequency was calculated by subtracting the average baseline firing one minute before distension. *P < 0.05, **P < 0.01, ***P < 0.001.

To determine the effect of galanin on LSN mechanosensitivity, we measured the effects of 500 nM galanin on the response to a ramp distension of the colon (0 – 80 mmHg) and observed a robust inhibition of spike frequency across the pressure range used, an effect that was most pronounced at the 80 mmHg (Fig. 1B and C). We therefore used a rapid phasic distension of the colon to noxious pressure (80 mmHg) to investigate the dose-dependent effect of galanin upon LSN mechanosensitivity. Galanin inhibited LSN firing in a dose-dependent manner, with the response to galanin being most marked on the third phasic distension after galanin infusion (subsequent values are therefore taken from this third distension, Fig. 1D). The maximal inhibition of the peak afferent response to phasic colonic distension was observed following the application of 500 nM galanin which produced an approximately 50% reduction in afferent mechanosensitivity (53.3 ± 12.8 spikes s^-1^ vs 25.2 ± 5.2 spikes s^-1^, N = 5, P < 0.0008, one- way ANOVA with Tukey’s test) and the IC_50_ calculated from the data was 65.7 nM (Fig. 1E). A similar inhibition by 500 nM galanin was observed of the sustained firing that occurred during the last 45 seconds of the distension (24.5 ± 8.6 spikes s^-1^ vs 9.3 ± 4.3 spikes s^-1^, N = 5, P = 0.013, unpaired t-test).

### Galanin inhibits noxious mechanically evoked neuronal excitation via GalR1

Having determined that galanin inhibited colonic afferent mechanosensitivity we next investigated the effect of GalR1 and GalR2 agonists on LSN activity. Using the same protocol as for galanin, the selective GalR1 agonist M617 (500 nM) elicited a reduction in LSN mechanosensitivity comparable in magnitude to galanin at 500 nM, such that the peak LSN response to mechanical distension was inhibited from 77.4 ± 10.3 spikes s^-1^ vs 46.5 ± 8.8 spikes s^-1^ (Fig. 2A; P = 0.0015, N = 4, paired t-test) and the sustained response was also inhibited from 37.3 ± 8.8 spikes s^-1^ vs 22.6 ± 6.9 spikes s^-1^ (Fig. 2B; P = 0.038, N = 4, paired t-test). The effect of 500 nM M617 was similar to that of 500 nM galanin: M671 attenuated peak firing frequency by 41.4 ± 7.5% and 500 nM galanin inhibited peak firing frequency by 49.3 ± 16.9% (M617 N = 4, galanin N = 5). By contrast, 1 µM of the endogenous GalR2 agonist spexin, which shows no binding to GalR1 (Kim et al., 2014), had no effect on LSN colonic afferent mechanosensitivity in response to phasic distension of the colon (Fig. 2C and D). These results show that galanin acts via GalR1 to inhibit LSN activity in response to distension of the colon.

**Figure 2.**
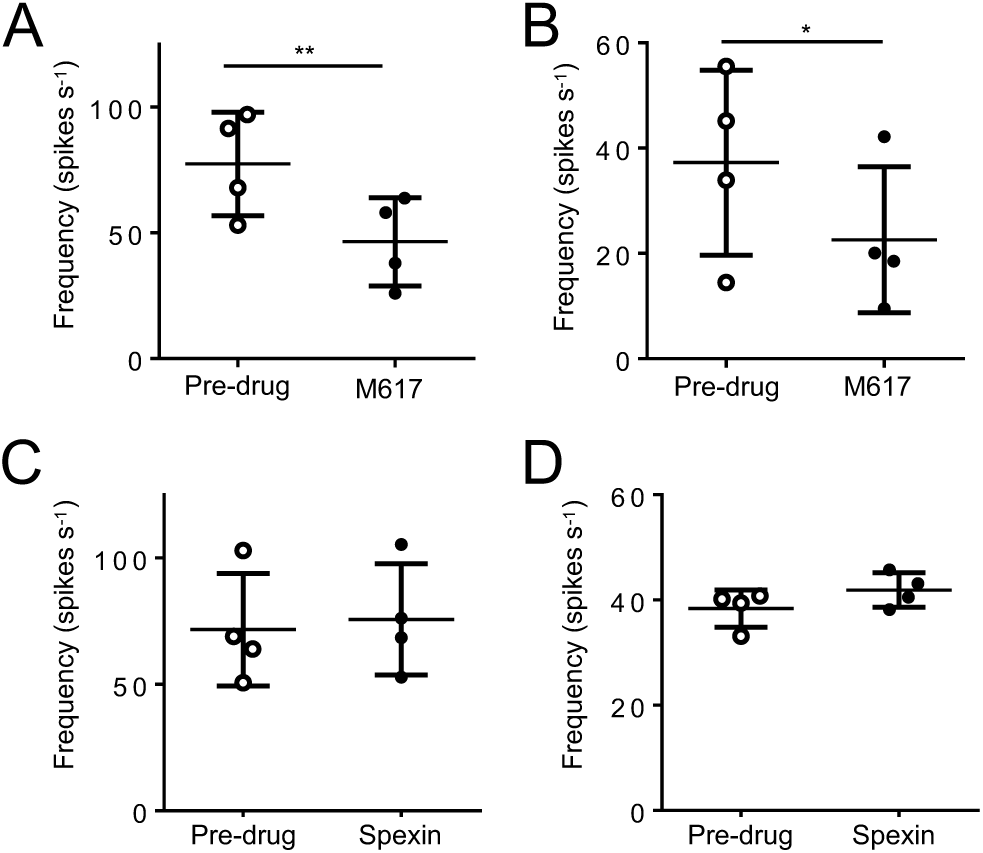
GalR1 activation inhibits LSN responses to phasic distension. (A) The GalR1 agonist M671 significantly attenuates both LSN peak (P = 0.0015, N = 4, paired t-test) and sustained (B; P = 0.0382, N = 4, paired t-test) firing in response to phasic distension of the colon to 80 mmHg. (C) The GalR2 agonist Spexin does not significantly attenuate or excite the peak (C; P = 0.245, N = 4, paired t-test) or sustained (D; P = 0.218, N = 4, paired t-test) response to phasic distension of the colon to 80 mmHg. Firing frequency was calculated by subtracting the average baseline firing one minute before distension. *P < 0.05, **P < 0.01

### Galanin inhibits mechanical hypersensitivity induced by inflammatory mediators

We next investigated the effect of galanin on the sensitisation of the LSN afferent response to colonic distension by intraluminal perfusion with an inflammatory soup (IS: ATP, histamine, PGE_2_, bradykinin, and serotonin) (Hockley et al., 2014; Su and Gebhart, 1998). The IS was intraluminally perfused following the third phasic distension (either alone or with 500 nM of galanin, Fig. 3A-D). As expected, IS significantly increased the peak firing frequency in response to phasic distension (31.2 ± 28.1 %, P < 0.05, N = 6, one-way ANOVA with Tukey’s test, Fig. 3E), whereas co-application of IS and galanin did not produce a significant change in peak afferent firing compared to control (Fig. 3E). A similar effect was observed with regard to the sustained firing frequency, such that IS application significantly increased sustained firing frequency (40.3 ± 26.7 %, P < 0.05, N = 6, one-way ANOVA with Tukey’s test), which was not observed when IS was combined with 500 nM galanin (Fig. 3F). Lastly, we also observed that the basal LSN activity increased by 81.6 ± 19.3% following IS application (P < 0.01, N = 6, one-way ANOVA with Tukey’s test), but that combination of IS and galanin did not significantly change the basal firing from the control group, suggesting again that the effects of galanin and IS cancel each other out (Fig. 3G).

**Figure 3.**
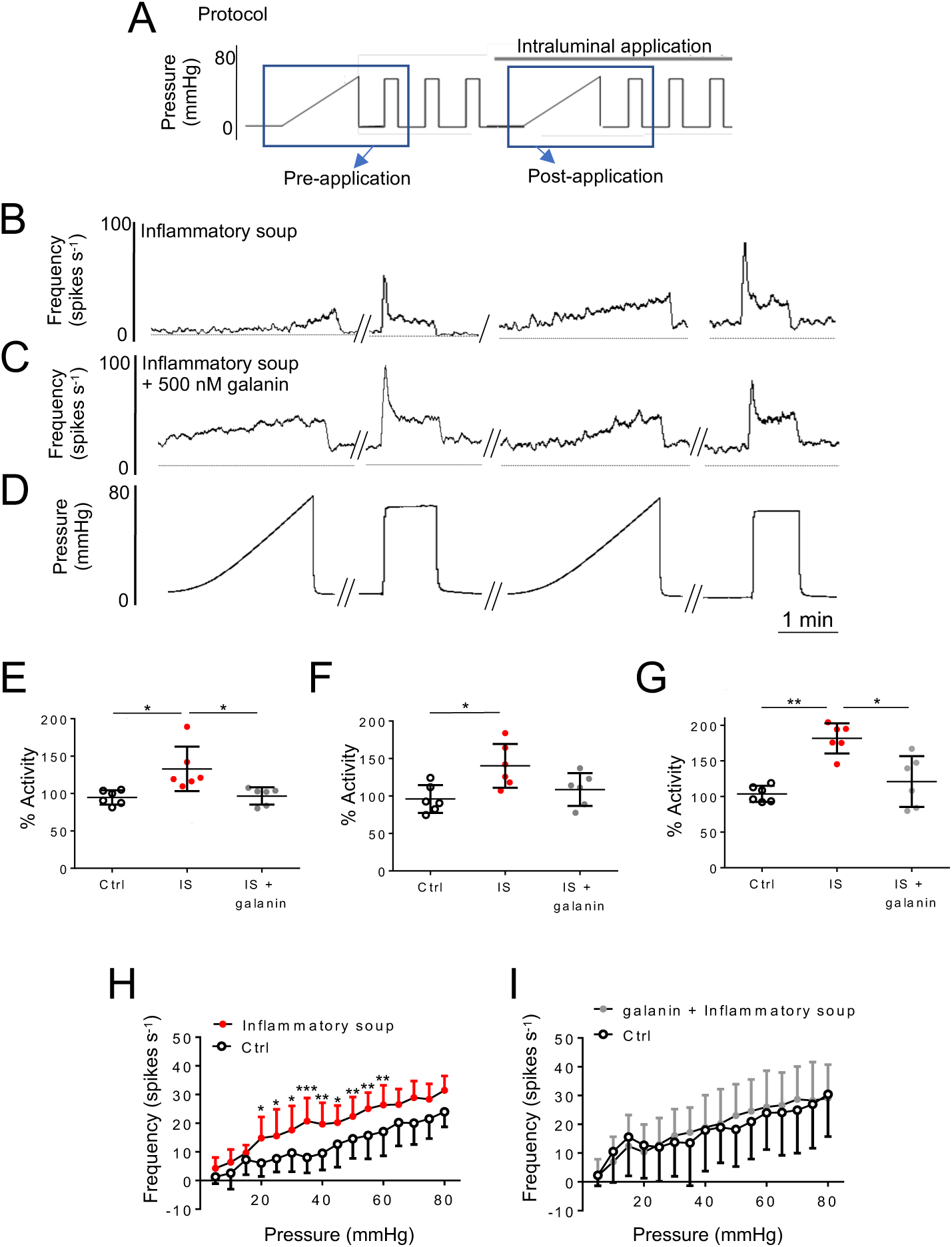
Galanin inhibits the effects of IS on LSN mechanosensitivity. (A) Schematic of pressure protocol used in the multi-unit recordings. Ramp and phasic distensions demonstrated below indicated with blue box, intraluminal application of drug indicated by black bar above the protocol. Example raw trace of ramp and phasic distension before and after the intraluminal application of inflammatory soup (B), a combination of Inflammatory soup and galanin (500 nM, C) and sample pressure distensions (D). Changes in peak (E), sustained (F) and basal (G) firing after intraluminal application of either inflammatory soup or a combination. Significant differences between groups tested by one- way ANOVA with Tukey’s test, *P < 0.05, **P < 0.01 (N = 6). Response profiles to ramp distension before and during intraluminal application of inflammatory soup (H) or a combination of IS and galanin (I, two-way repeated-measures ANOVA with Sidak’s post-hoc test, N = 6). Firing frequency was calculated by subtracting the average baseline firing one minute before distension. Significance indicated by *P < 0.05, **P < 0.01, ***P< 0.001.

To further investigate LSN hypersensitivity caused by IS application and how this is affected by galanin, we used a ramp distention (0 to 80 mmHg over ∼2.5 minutes). As expected, IS significantly increased activity across a broad range of pressures (Fig. 3H, N = 6), and again the combination of IS and galanin resulted in a pressure-induced activity relationship that was not significantly different to control (Fig. 3I, N = 6). These results demonstrate that galanin inhibits inflammatory mediator induced mechanical hypersensitivity of LSN afferents in the colon.

### Galanin does not inhibit DSS-induced mechanical hypersensitivity

Using the DSS model of colitis, we observed that DSS treated mice showed significant weight loss from day 4 onwards compared to untreated mice (e.g. on day 7, 82.1 ± 1.9% vs 103.9 ± 3.6 % of starting weight, N = 10, P <0.0001, unpaired t-test) (Fig. 4A), and also a significant increase in disease activity index (DAI, Fig. 4B). Following dissection, colon length was observed to be significantly decreased (Fig. 4C and D) and the colon wet weight to length ratio was also significantly increased in DSS mice compared to the control group (Fig. 4E). We observed both significant macroscopic and histological damage (Fig. 4F-H) in colon sections from DSS mice compared to those from healthy mice, as well as significant thickening of the muscular layer of the colon in the DSS group (Fig. 4I) as others have observed (Marrero et al., 2000; Sánchez-Fidalgo et al., 2012). Lastly, the extracellular matrix polysaccharide hyaluronan (HA) was largely absent from the colon epithelium of DSS treated mice further demonstrating a breakdown in tissue integrity (Fig. 4J).

**Figure 4.**
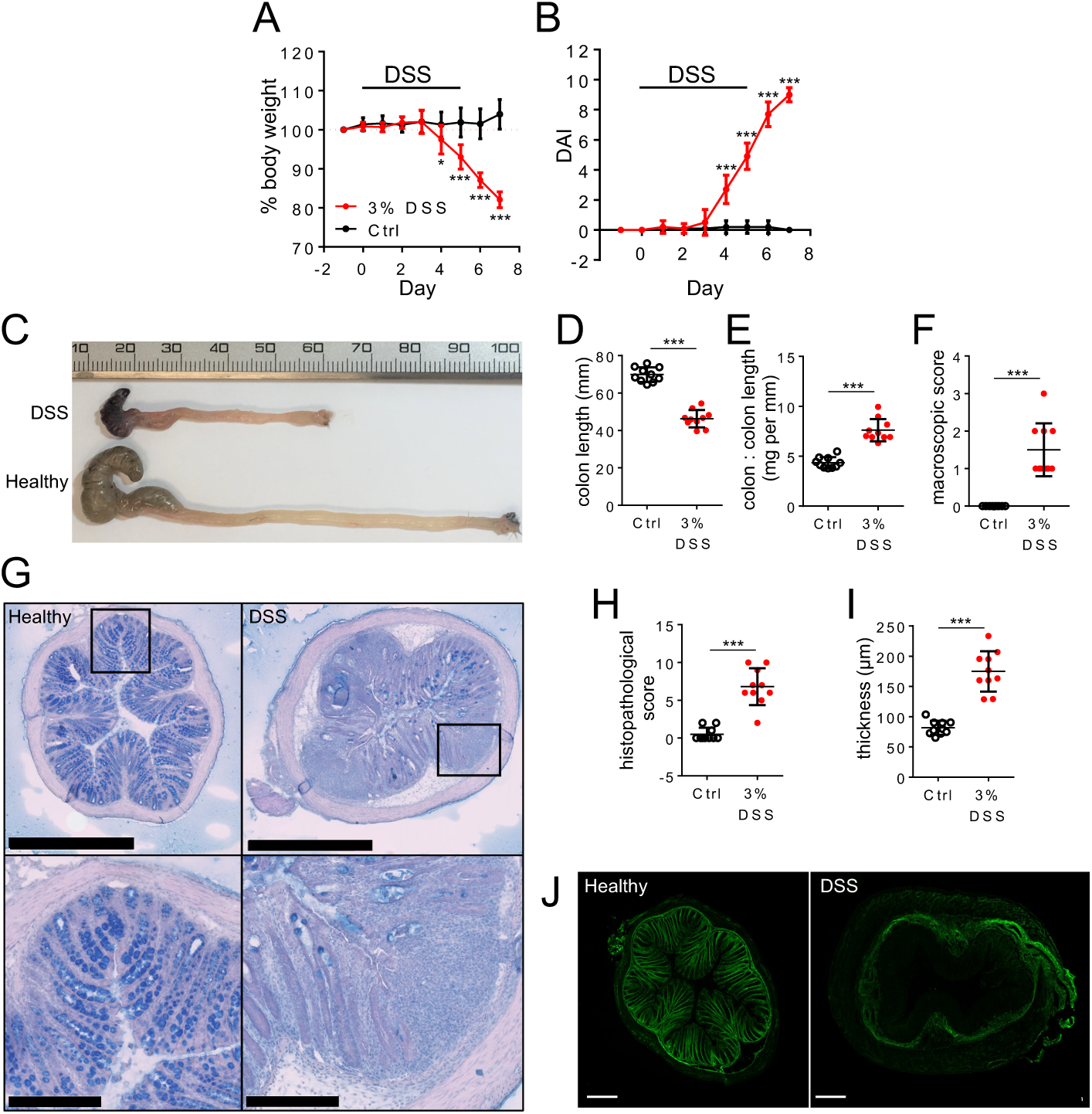
DSS induces weight loss and macroscopic changes to colon histology. (A) Body weight of DSS treated mice was significantly reduced compared to untreated controls (* P < 0.05, *** P < 0.001, N = 10, two-way repeated-measures ANOVA with Sidak’s post-hoc). (B) DAI significantly increases in colitis-induced mice (*** = P < 0.001, N = 10, two-way repeated-measures ANOVA with Sidak’s post-hoc test). DAI = disease activity index; assessment of inflammation by clinical parameters. (C) Photograph of healthy and colitis-induced colons from caecum (left side) to anus (right side). (D) Colon length is significantly reduced in DSS mice (P < 0.001, N = 10, paired t-test). (E) Colon weight to length ratio significantly increased in DSS mice (P < 0.001, N = 10, paired t-test). (F) Macroscopic score is based on visual assessment of ulceration and hyperaemia of the colon (P < 0.001, N = 10, paired t-test). (G) H&E with alcian blue staining of colonic tissue. In DSS mice, there was active inflammation, and crypt or surface epithelial damage compared to untreated controls. Areas defined by black boxes are magnified in the lower images; scale bar for top images is 1 mm and for bottom images 250 μm. (H) Histology score significantly increases in DSS mice (P < 0.001, N = 10, paired t-test) and the colonic muscle layer becomes significantly thicker in DSS mice (I, P < 0.001, N = 10, paired t-test). (J) Changes in hyaluronic acid binding protein (HABP; green) arrangement and distribution in colitis-induced and healthy colons.

Using the DSS model, we investigated if the inhibition of LSN activity by galanin in healthy mice (Figure 1) was maintained in LSN from mice undergoing DSS-induced colonic inflammation. Using a ramp distention, we observed that the LSN isolated from DSS treated mice produced a greater response at non-nociceptive pressure (20 mmHg, 6.9 ± 5 spikes s^-1^ vs 19.8 ± 4.7 spikes s^-1^, P = 0.0184, N = 6, t-test), but not at nociceptive pressure compared to the LSN from healthy mice (34.7 ± 8.1 spikes s^-1^ vs 39.7 ± 3.2 spikes s^-^1, P = 0.137, N = 6, Student’s T-test; 80 mmHg, Fig. 5A). In addition, the basal firing of LSN from DSS treated mice was significantly greater than that of healthy mice (3.9 ± 1.4 spikes s^-1^ vs 28.4 ± 13.4 spikes s^-1^, P = 0.012, N = 6, Student’s T-test). Together with the ramp distension response data, these results suggest that DSS induced a state of visceral hypersensitivity at the level of the primary afferent. Whereas galanin was observed to significantly inhibit the peak and sustained LSN responses induced by a phasic distension to 80 mmHg of the colon in healthy mice (Fig. 1), no such inhibitory effect was observed when measuring LSN activity in nerves isolated from DSS mice: peak firing frequency (65.7 ± 16.6 spikes s^-1^ vs 66.6 ± 22.3 spikes s^-1^, P = 0.81, N = 6, paired t-test; Fig. 5B) and sustained firing frequency (34.1 ± 9.9 spikes s^-1^ vs 41.9 ± 14 spikes s^-1^, P = 0.54, N = 6, paired t-test; Fig. 5C) induced by a phasic distension to 80 mmHg were affected by administration of galanin. These results indicate that the inhibitory action of galanin is lost in tissue isolated from mice with acute colitis.

**Figure 5.**
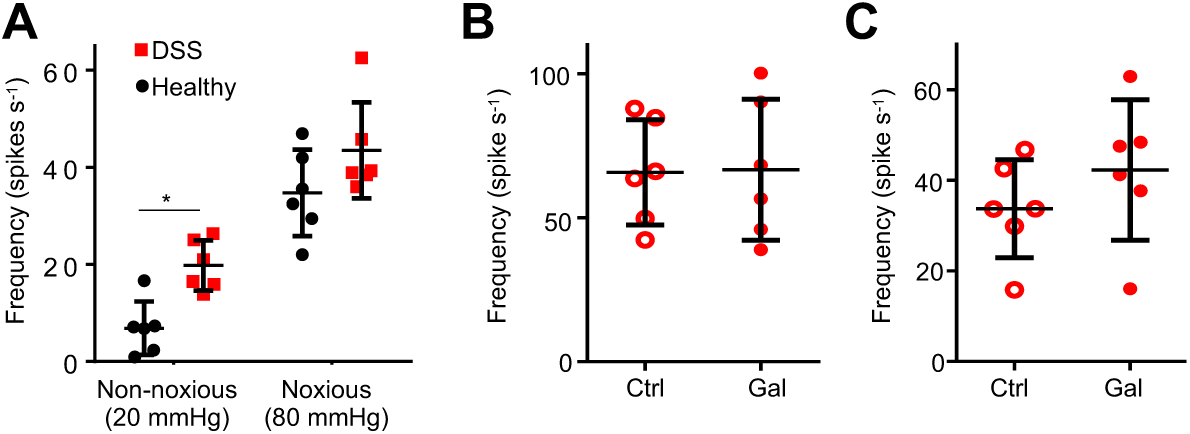
Galanin does not inhibit DSS-induced LSN hypersensitivity. (A) DSS increases the response to non-noxious (20 mmHg), but not noxious (80 mmHg) pressures using a ramp distension protocol; Student’s t-test between groups * P < 0.05. 500 nM galanin does not inhibit peak (B) or sustained (C) responses to phasic distension of the colon to 80 mmHg in colons isolated from DSS treated mice (paired t-test, N = 6). Firing frequency was calculated by subtracting the average baseline firing one minute before distension.

## Discussion

Our data demonstrate that galanin inhibits LSN responses to noxious mechanical stimuli and the sensitisation of colonic mechanosensitivity by acute application of inflammatory mediators. This inhibitory effect of galanin not seen in tissue isolated from mice with colitis. Our data further suggest that GalR1 mediates the inhibitory effect of galanin. These data fit alongside galanin’s reported function in the upper GI tract (Page et al., 2005) and parallels with the function of galanin in somatic sensory innervation (Flatters et al., 2003; Heppelmann et al., 2000).

Galanin is expressed by multiple cell types in the distal colon including enterochromaffin cells and fibroblasts (Schäfermeyer et al., 2004; Yamamoto et al., 2014), and, as we show here, a subset of colonic sensory neurones also express galanin. Galanin has been implicated as a modulator of numerous activities in the GI tract including regulation of neurotransmitter release, motility and secretion (Benya et al., 1999; Sternini et al., 2004). With regard to the transmission of sensory information from the GI tract, galanin has been shown to modulate mechanosensitivity of gastro-oesophageal vagal afferents with predominantly inhibitory actions on individual fibres via GalR1 (Page et al., 2005; Page et al., 2007). Our data build on this by demonstrating that galanin inhibits LSN mechanosensitivity and likely does so though GalR1 activation.

Mechanical distension of the colon is capable of producing pain in humans and nociceptive behaviour in animals (Gebhart, 2004; Ness et al., 1990). Consequently, distension is a useful tool when defining noxious responses in afferent preparations and examining hyperexcitability to colonic inflammation. We find from multi-unit recordings of LSN activity that galanin inhibits mechanically-evoked responses to distension of the distal colon to noxious pressures and the sensitisation of these responses following addition of inflammatory stimuli. This suggests that galanin receptors, specifically in the case of the LSN GalR1, function as integral regulators of neuronal excitability. Multiple populations of sensory afferents innervate the distal colon with differing sensitivities to mechanical stimuli (e.g. stretch, stroke and von Frey hair probing of their receptive fields) (Brierley et al., 2004) and whilst we did not seek to characterise these groups more specifically, we did observe inhibitory effects of galanin across the full range of distension pressures from physiological through to noxious (i.e. 0 – 80 mmHg, Fig. 1C). When examining the expression of galanin, both colonic sensory neurone RNA-sequencing and immunohistochemistry (Fig. 1A) demonstrate that galanin is expressed in putative nociceptors (Hockley et al., 2019). With regard to GalR1, colonic sensory neurone RNA-sequencing indicates that it is predominantly expressed in neurones expressing nociceptor markers, such as Trpv1 and Gfrα3, as well as in a population of neurones expressing the mechanosensitive ion channel Piezo2, an expression pattern that likely explains the inhibitory impact of galanin on LSN activity across a wide range of pressures (Fig. 1C). The GalR1 receptor is coupled to G protein-coupled inwardly rectifying potassium channels (GIRKs) giving rise to hyperpolarization (Walker et al., 1997; Smith et al., 1998), an effect that would account for the inhibitory activity of galanin observed in this study. This conclusion is further supported by data demonstrating that inhibition of LSN activity was also produced by the GalR1 agonist M617, but not the GalR2 agonist spexin (Fig. 2).

As observed in previous studies (Su and Gebhart, 1998), we found that intraluminal application of IS produced robust LSN hyperexcitability to mechanical stimuli (Fig. 3). Part of this hyperexcitability likely results from recruitment of ‘silent’ or mechanically insensitive afferents in both the LSN and PN (Feng and Gebhart, 2011), potentially through disinhibition of Piezo2 (Prato et al., 2017). The ability of galanin to reverse the mechanically hypersensitivity induced by the IS correlates with the fact that a variety of inflammatory mediator receptors are present in colonic sensory neurones that also express GalR1 (Hockley et al., 2019). However, by contrast, galanin was not able to reduce the mechanical hypersensitivity present in LSN isolated from mice treated with DSS to induce a state of colitis.

Why is it that galanin counteracts the acute effects of inflammatory mediators on LSN activity, but has no such effect on LSN activity in a mouse model of colitis? It has been observed that galanin receptor expression is altered in certain models of inflammation and hence differential receptor expression could lead to galanin no longer exerting an inhibitors effect. For example, following hind-paw injection of carrageenan in rats, GalR1 mRNA expression in DRG neurones (L4 and L5) decreases (Xu et al., 1996) and a similar decrease in GalR1 expression could occur in the DSS model and hence galanin is no longer able to inhibit mechanically-evoked LSN activity; alternatively, there could be an increase in the expression of excitatory GalR2. The absence of reliable tools to investigate GalR protein levels and the validity of current antibodies being uncertain (Lu and Bartfai, 2009) makes quantifying the protein level of GalRs in colonic afferents a significant challenge. However, there are also alternative explanations for the lack of measurable galanin activity. For example, multiple inflammatory mediators are released from inflamed colon tissue (Spiller and Major, 2016), which would act upon a broad range of afferents to induce a variety of transcriptional changes and post-translational modifications, i.e. both GalR1+ve and GalR1-ve afferents, and thus the mechanical hypersensitivity observed in LSN isolated from DSS treated mice is likely at least partially mediated via GalR1-ve afferents. Therefore, any effect of galanin on the whole-nerve response may simply be overcome by the overall level of sensitisation. A further explanation would be that the coupling of GalR1 is altered in inflammation due to altered expression of G proteins and/or that the signalling of galanin at GalR1 becomes biased towards different pathways.

In conclusion, we have shown a hitherto unreported role for galanin in the modulation of LSN function in the distal colon: galanin inhibits LSN mechanosensitivity and acute mechanical hypersensitivity induced by an IS. Future work should elucidate mechanisms underpinning why galanin is unable to exert any inhibition on LSN mechanical hypersensitivity following prolonged (*in vivo*) inflammation. Nevertheless, our data suggest that targeting GalR1 could provide a new route to treating visceral pain under specific circumstances.

## Additional information

### Author contributions

TST, JH and ESS conceived and designed the research. TST performed the experiments and analysed the data with assistance from PK and SJ. TST drafted the manuscript and all authors edited, revised and approved the final version of the manuscript submitted for publication. All authors agree to be accountable for all aspects of the work in ensuring that questions related to the accuracy or integrity of any part of the work are appropriately investigated and resolved. All persons designated as authors qualify for authorship, and all those who qualify for authorship are listed.

